# Wound healing and angiogenic profiling of dermal endothelial cells isolated from people with type 2 diabetes

**DOI:** 10.1101/2025.07.18.665569

**Authors:** James Shadiow, Corey E. Mazo, Pallavi Varshney, Jeongjin J. Kim, Alexander Ahn, Crystal M. Holmes, Michael E. Munson, Brian M. Schmidt, Sascha N. Goonewardena, Richard D. Minshall, Thomas P.J. Solomon, Andrew T. Ludlow, Jacob M. Haus

## Abstract

Impaired wound healing in type 2 diabetes (T2D) is associated with microvascular dysfunction and remains a significant clinical challenge. We aimed to determine whether primary human dermal microvascular endothelial cells (HDMVECs) from individuals with T2D exhibit abnormal cellular functions, and whether exposure to T2D serum impacts healthy endothelial function. In Experiment 1, T2D-HDMVECs displayed paradoxically higher migratory and angiogenic capacities than their healthy counterparts, despite markedly reduced eNOS expression and disrupted endothelial-identity gene expression. In Experiments 2 and 3, healthy HDMVECs showed decreased tube formation, nitric oxide production, and Notch/angiogenesis-related gene expression after exposure to both healthy and T2D serum, suggesting the presence of serum-derived factors that inhibit these pathways. However, T2D-HDMVECs remained largely unresponsive to these serum-driven effects, reinforcing an “intrinsic” reprogramming of T2D endothelial cells. Overall, our data reveal a complex interplay between cell-autonomous alterations and extrinsic signals in diabetic endothelial dysfunction. Therapeutic strategies targeting both intrinsic cellular programs (e.g., eNOS, Notch signaling) and the circulating milieu may represent promising avenues for enhancing wound repair in patients with T2D.

## Introduction

Impaired wound healing in type 2 diabetes (T2D) is a leading cause of lower-extremity amputation and is associated with increased mortality [1, 2]. Multiple factors delay wound healing, including reduced angiogenesis and impaired microvascular perfusion, reducing oxygen delivery to the affected tissue [2-5]. However, the pathophysiological processes are not fully understood.

Angiogenesis is essential for tissue repair and involves growth factors and cytokines that trigger endothelial cell migration and proliferation to stimulate tube formation and maturation of new blood vessels [3, 6, 7]. However, in T2D, pathological angiogenesis can occur, whereby excessive or abnormal endothelial cell outgrowth leads to pathological blood vessel formation and ineffective microvasculature, impairing wound healing [8, 9]. Similarly, microvascular perfusion is essential for delivering nutrients and oxygen to the wound site to facilitate healing [6, 8], but this function is impaired in T2D [10-12].

Despite evidence that reduced angiogenesis and microvascular perfusion are key mediators of impaired wound healing in T2D, the precise pathophysiology is unclear because intrinsic alterations in cellular programming or cellular exposure to the diabetic milieu could drive them [6, 7, 10, 12]. One such intrinsic alteration involves the loss of endothelial identity via endothelial-to-mesenchymal transition (EndMT), which contributes to the development of severe microvascular complications [13, 14] and is promoted by several factors, including TGF-β1, inflammatory cytokines, oxidative stress, hyperglycemia, hypoxia, and shear stress [15-17]. Many of these factors are also characteristics of the T2D phenotype.

A major regulator of angiogenesis and endothelial function is nitric oxide. We have previously shown that endothelial nitric oxide synthase (eNOS) activity plays a key role in chronic inflammation, angiogenesis, and wound healing in endothelial cells [18, 19] and in a mouse model of obesity-induced T2D [19, 20]. However, clinical translation is needed, and the specific role of eNOS in the behavior of human endothelial cells derived from individuals with T2D remains incompletely defined.

Another promising target is Notch signaling, which regulates cellular processes essential for wound healing, such as angiogenesis [21, 22]. Studies in rodents have found that activation or inhibition of Notch signaling can alter the behavior of cultured vascular endothelial cells [23-25]. Evidence also indicates that Notch signaling is differentially regulated in rodent models of T2D in various tissues, including the kidney and the intestine [26, 27], and can regulate injury repair in bone [24]. Consequently, Notch pathways have been identified as a potential therapeutic target in diabetic foot ulcers [22]. However, specific studies directly linking Notch signaling to diabetic wound healing are limited.

The circulating environment in T2D contains high levels of glucose, cytokines, and chemokines that directly affect vascular endothelial cells and can induce vascular leakage by activating coagulation factors, such as thrombin [28]. Despite extensive studies on extrinsic factors such as high glucose and TGF-β1, effective treatments that correct environmental abnormalities are lacking. Indeed, even when blood glucose levels are normalized, people with T2D can continue to exhibit vascular complications [29, 30], suggesting that intrinsic alterations in cellular programming persist when the exposure to the diabetic milieu is removed. These findings underscore the interplay between environmental influences and endothelial phenotypes, driving diabetic vasculopathy.

In summary, it is unclear whether impaired wound healing in T2D is caused by an inherent cellular defect or cellular exposure to the diabetic milieu. Therefore, we aimed to determine whether circulating factors in people with T2D affect wound healing in endothelial cells, ultimately aiming to identify potential therapeutic targets to enhance wound repair in patients with T2D-related foot ulcers. We hypothesized that primary human dermal microvascular endothelial cells (HDMVECs) from people with T2D would exhibit abnormal endothelial cell function (impaired migration and tube formation). We further hypothesized that exposing HDMVECs collected from healthy individuals to serum collected from people with T2D would impair endothelial cell function via a Notch-related mechanism.

## Methods

### Experimental Design

Firstly, to model wound healing, we used a scratch assay to determine whether microvascular endothelial cells collected from people with T2D exhibited abnormal repair compared to cells collected from people without known health conditions. Then, to determine whether the diabetic milieu blunted repair, we incubated “healthy” cells with sera collected from people with TDM, and vice versa.

### Participants and blood sampling

All study protocols were ethically approved by the Institutional Review Boards of the University of Michigan and performed according to the Declaration of Helsinki. People with T2D (n=11) and people without known health conditions (n=10) were recruited from the local area (Ann Arbor, MI) and underwent medical screening. Males and females of all races aged 18-65 years were eligible for inclusion and recruited to a healthy or T2D group. Inclusion to the T2D group required a clinical diagnosis of T2D confirmed by a 75 g oral glucose tolerance test (OGTT) [31] and medical management of T2D by metformin alone. Individuals were excluded if they had undergone greater than 2 kg weight change in the last 6 months; had existing cardiovascular, cerebrovascular, renal, or hematological disease, cancer, or other diseases suspected to impact study outcomes; currently used tobacco or nicotine; were pregnant/lactating; or taking medications for T2D management other than metformin. People with impaired fasting glucose or impaired glucose tolerance, as measured by OGTT, were also excluded.

Participants who met the inclusion criteria provided informed consent for future sample use and were invited to a blood sample collection visit in the morning following an overnight fast of 10-12 hours. Participants were instructed to refrain from consuming alcohol (for 48 hours) and caffeine (for 24 hours) before the visit and to avoid exercising for at least 24 hours. Participants were also asked to refrain from taking medications and supplements known to influence the primary outcomes on the morning of blood sampling (e.g., anti-hypertensives, statins). Participants arrived in the laboratory at 8 am, and 10 ml of venous blood was obtained from an antecubital vein and collected into serum separator tubes. The blood was centrifuged, and the serum was separated and stored at -80°C. On the day of the experiments described below, serum samples were thawed and pooled before use in cell culture. These pools are referred to as Healthy-serum and T2D-serum.

### Primary human dermal microvascular endothelial cell culture

Primary HDMVECs were obtained from people without known health conditions (H-6064, lot: SMN0519017, Cell Biologics, Chicago, IL) and people with T2D (HD2-6064, lot: F112817Y50IIAM, Cell Biologics). These are referred to as Healthy-HDMVECs and T2D-HDMVECs, respectively. Cells from passages 5-9 were propagated in T-75 flasks (at least 10×10^3^ cells/cm^2^) and maintained in serum-free endothelial growth medium (EGM-2; #CC-3162, Lonza, Walkersville, MD) at 37°C and 5% CO_2_ throughout the experiments. The growth medium was replaced every other day until cells reached 70% confluence, after which the medium was changed daily. Three experiments were completed using cells grown and maintained in EGM-2 at >90% confluency:

***Experiment 1***. N=12 experimental replicates were completed for each cell type. Healthy-HDMVECs and T2D-HDMVECs were grown to 90% confluency, and a 200 μL pipette tip was used to scratch a smooth line vertically in the center of the plate (the scratch assay). The plate was washed, fresh EGM-2 media was added, and the cells were incubated for a further 24 hours. Scratch wounds were imaged before and after the 24-hour repair period to examine cell migration. The cells were then washed, collected, and used to assess tube formation, gene expression of Nos3, Cdh5, Cav1, Acta2, and protein expression of eNOS.

***Experiment 2***. N=6 experimental replicates were completed for each cell type. Healthy-HDMVECs and T2D-HDMVECs were grown to 90% confluency, the plate was scratched as described above, washed, and exposed to one of three conditions: (i) serum-free EGM-2 media (referred to as *No Treatment, NT*), (ii) EGM-2 media supplemented with 30% Healthy-serum, (iii) EGM-2 media supplemented with 30% T2D-serum, as described previously [32, 33]. Cells were incubated for 24 hours under these conditions, then washed and serum-starved for 2 hours in EBM-2. Before and after this period, cell migration was assessed, and conditioned media samples were collected to measure nitrite production. A treatment period of 24 hours was chosen based on our unpublished data supporting the greatest effects of treatment on cellular functions. Finally, the cells were washed, collected, and used to assess tube formation. An overview of this study design is shown in **Fig. 1**.

**Fig. 1.**
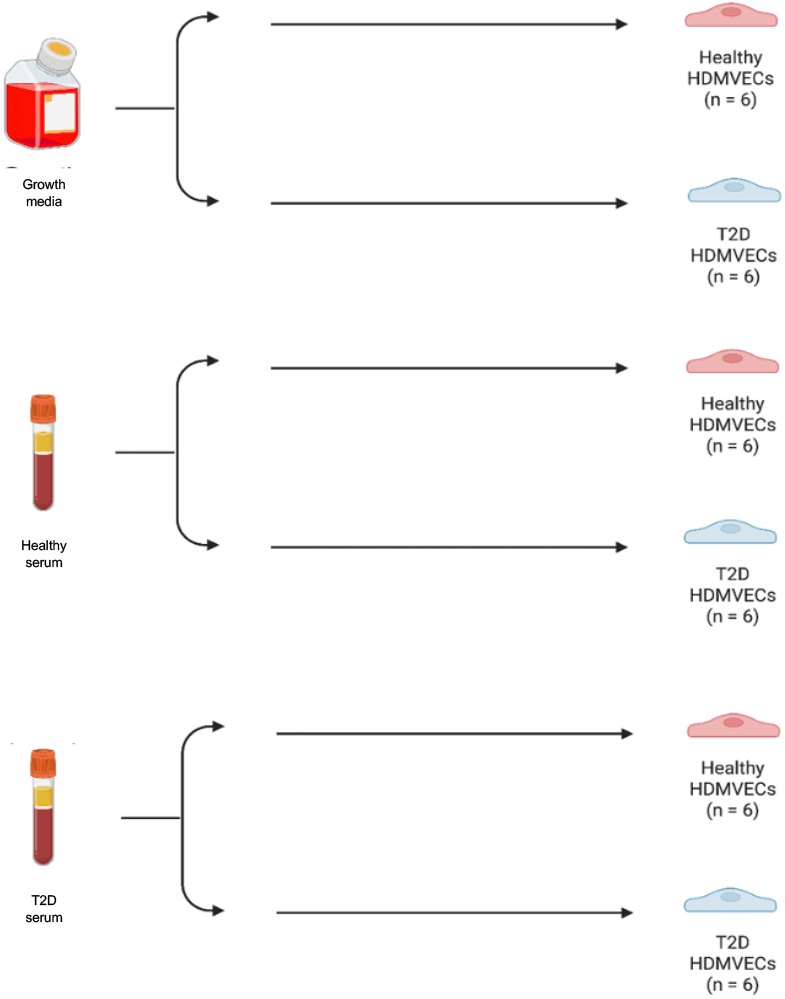
Experimental design for the pooled serum treatments of Healthy-HDMVECs and T2D-HDMVECs. HDMVECs isolated from people without known health conditions (Healthy HDMVECs) and people with type 2 diabetes (T2D HDMVECs) were incubated in either growth media, serum collected from people without known health conditions (Healthy-serum), or serum collected from people with type 2 diabetes (T2D-serum).

***Experiment 3***. N=6 experimental replicates were completed for each cell type. The procedures in Experiment 2 were repeated; however, instead of measuring cell function and nitrite at the end of the treatments, cells were washed, collected, and used to assess gene expression of Hes1, Hey1, Dll4, Jag1, Vegfr2, Notch1, Notch2, and Notch3.

### Cell migration

Cells were imaged via Keyence All-in-One BZ-X microscope (Keyence, Itasca, IL) at baseline and 24 hours post-scratch. Cell migration was analyzed using the Wound Healing Size Tool, a plug-in for ImageJ (v.2.9.0/1.53t) [34]. A migration index, which assesses the total scratch area closure, was calculated as:

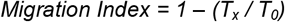

### Tube formation

Following the treatments described above, cells were counted using a Countess II FL Automated Cell Counter (ThermoFisher) then seeded in triplicate at 10×10^4^ cells/well in 96-well plates preloaded with Matrigel matrix (50 µL/well; 356234, Corning, Glendale, AZ). Plates were incubated for 24 hours and then imaged using an EVOS FL microscope (ThermoFisher). The Angiogenesis Analyzer plug-in for ImageJ was used to analyze tube formation [35].

### Gene expression

RNA was extracted from cells with RNeasy Plus Universal Kit (73404, Qiagen) according to the manufacturer’s instructions. RNA was quantified via NanoDrop One^c^ spectrophotometer (ThermoFisher). RNA (80–150 ng) was reverse transcribed using iScript Advanced reverse transcriptase kit (BioRad) using the manufacturer’s protocol. Confirmation of complementary DNA (cDNA) library creation protocol was assessed via agarose gel probing for *Gapdh*. Droplet digital PCR (ddPCR) was used to quantify transcripts providing an absolute count of target cDNA copies of *Acta2, Cav1, Cdh5, Dll4, Hes1, Hey1, Jag1, Nos3, Notch1, Notch2, Notch3,* and *Vegfr2*, as previously described [36]. Primer sequences are provided in **Table 1,** and the primers were validated via polyacrylamide gel electrophoresis against non-template, negative controls in which the cDNA was substituted for nuclease-free water. In brief, ddPCR was performed by combining cDNA with QX200 ddPCR EvaGreen Supermix (1864034, BioRad), primers, and nuclease-free water. Non-template, negative controls were included alongside each of the assays. Droplets were generated with a QX200 Droplet Generator (1864002, BioRad). The resultant droplet suspension then underwent 40 cycles of PCR and was analyzed using a QX200 Droplet Reader (1864003, BioRad) by counting the droplets positive for fluorescence. Transcript data are presented as the copy number per 1 ng of total RNA input.

**Table 1.**
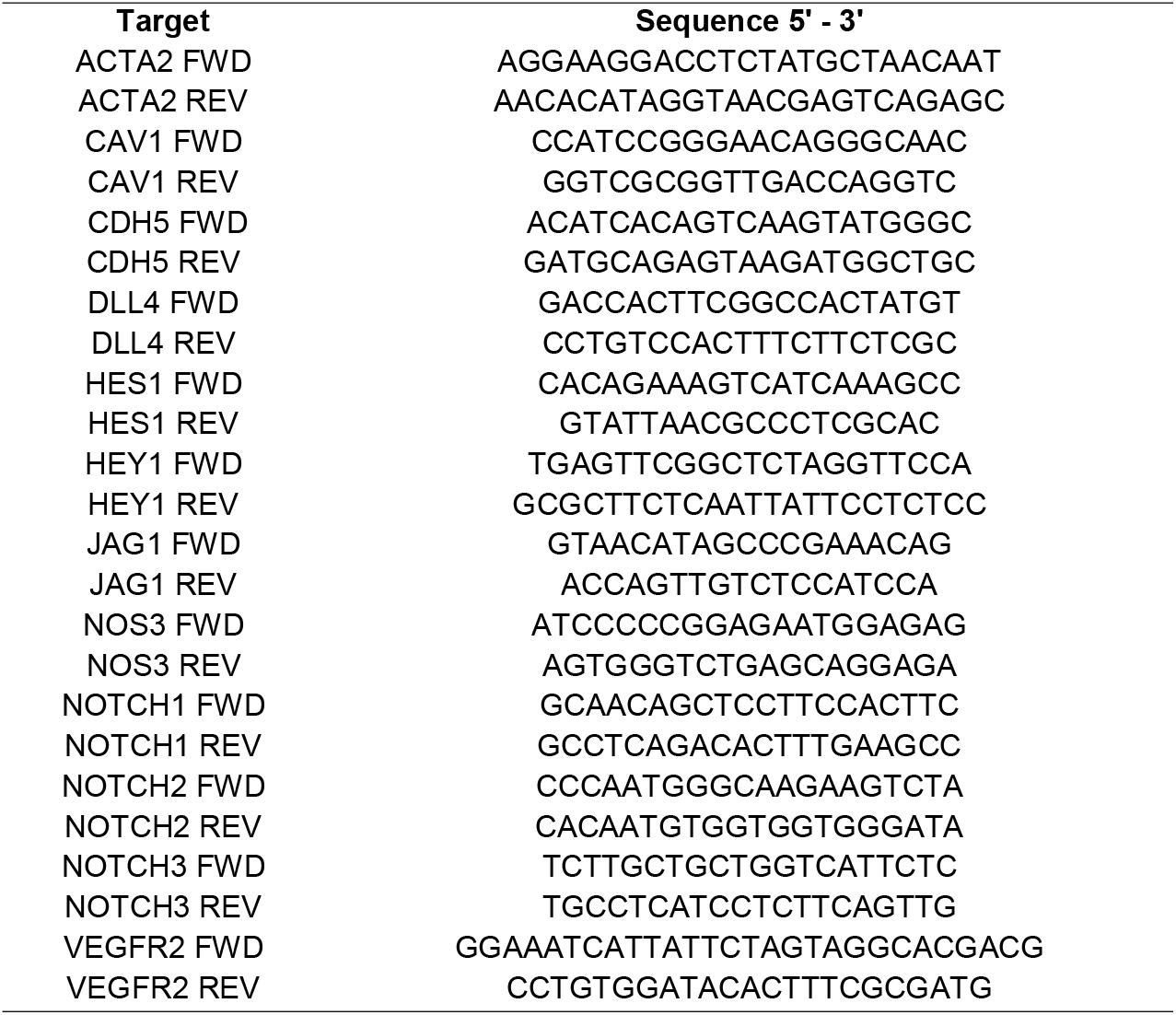
Primer sequences.

### Protein expression

To preserve protein phosphorylation status at the time of collection, cells were lysed in RIPA buffer (ThermoFisher) supplemented with 1X protease/phosphatase inhibitor cocktail (5872, Cell Signaling Technology, Danvers, MA). Protein concentrations were assessed using a Pierce 660nm protein assay reagent (22660, ThermoFisher) read using a NanoDrop One^c^ spectrophotometer (ND-ONE-W, ThermoFisher). Cell lysates containing equal amounts of total protein were diluted in 4X Laemmli Buffer (1610747, BioRad, Hercules, CA) with 5% β-mercaptoethanol (1610710, BioRad) before heating at 90°C for 10 min and separating on 4-20% pre-cast Criterion TGX gels (5671093, BioRad). Separated proteins were then transferred to nitrocellulose membranes via Transblot Turbo (BioRad, Lincoln, NE) and blocked with Protein-Free Blocking Buffer (PFBB, 92780003, Li-Cor, Lincoln, NE) for 1 hour at room temperature. Membranes were probed with primary antibody against eNOS (1:750, 5880, Cell Signaling Technology) diluted in PFBB + 0.1% Tween-20 overnight at 4°C. Membranes were then probed with 800 fluorophore-conjugated anti-mouse secondary antibody (1:20,000, Li-Cor, Lincoln, NE) diluted in PFBB + 0.1% Tween-20 for 1 hour at room temperature. Membranes were imaged using Odyssey CLx Imaging System (Li-Cor). Membranes were then rinsed in ultrapure water prior to incubating in REVERT total protein stain (926-11011, Li-Cor) for 5 minutes at room temperature and rinsed in REVERT wash solution before Odyssey CLx (Li-Cor) imaging to capture signal for total protein. Protein fluorescence was quantified using the WB quantification function in Image Studio software (V4.0.21; Li-Cor). Total protein was quantified using EmpiriaStudio software’s total protein quantification function (V2.0.0.131; Li-Cor). eNOS protein expression was normalized to total protein.

### Nitrite

Nitrite concentrations in conditioned media were measured in duplicate using a Griess reaction assay (Cayman Chemicals, Ann Arbor, MI). Nitrite production was calculated as the difference in nitrite concentrations in cell supernatants between the post- and pre-treatment conditions.

### Statistical analysis

All data are presented as mean ± standard error (SEM). Statistical analyses were performed using Prism 9.0. When comparing T2D vs. healthy HDMVECs, Shapiro-Wilk tests were used to test for distribution of data, and independent t-tests or Mann-Whitney U tests were used to identify group differences. When comparing T2D vs. healthy HDMVECs under multiple conditions, a two-way ANOVA was used with Šidák’s correction to adjust for multiple comparisons. Statistical significance was determined if *p*<0.05.

## Results

### Phenotypic differences between Healthy-HDMVECs and T2D-HDMVECs

T2D-HDMVECs displayed an elongated and disorganized cell morphology compared to Healthy-HDMVECs (**Fig. 2A and Fig. 2B**). The migration index (i.e., gap closure) following the plate scratch (injury) was 52% greater in T2D-HDMVECs than Healthy-HDMVECs (*p* < 0.001; **Fig. 2A**). Tube formation (i.e., angiogenesis) was also elevated in T2D-HDMVECs, which exhibited a 38% greater network length (*p* = 0.055) and 186% greater number of branches, compared to Healthy-HDMVECs (*p* < 0.001; **Fig. 2B**). Meanwhile, eNOS protein expression was 90% lower in T2D-HDMVECs (*p* = 0.004; **Fig. 2C**) coinciding with a markedly reduced *Nos3* gene expression, compared to Healthy-HDMVECs (*p* < 0.001; **Fig. 2D**). Furthermore, T2D-HDMVECs exhibited 89% lower *Cdh5* and 82% lower *Cav1* gene expression compared to Healthy-HDMVECs (*p* < 0.001, *p* = 0.002, respectively; **Fig. 2D**), with no difference in *Acta2* expression (P>0.05; **Fig. 2D**).

**Fig. 2.**
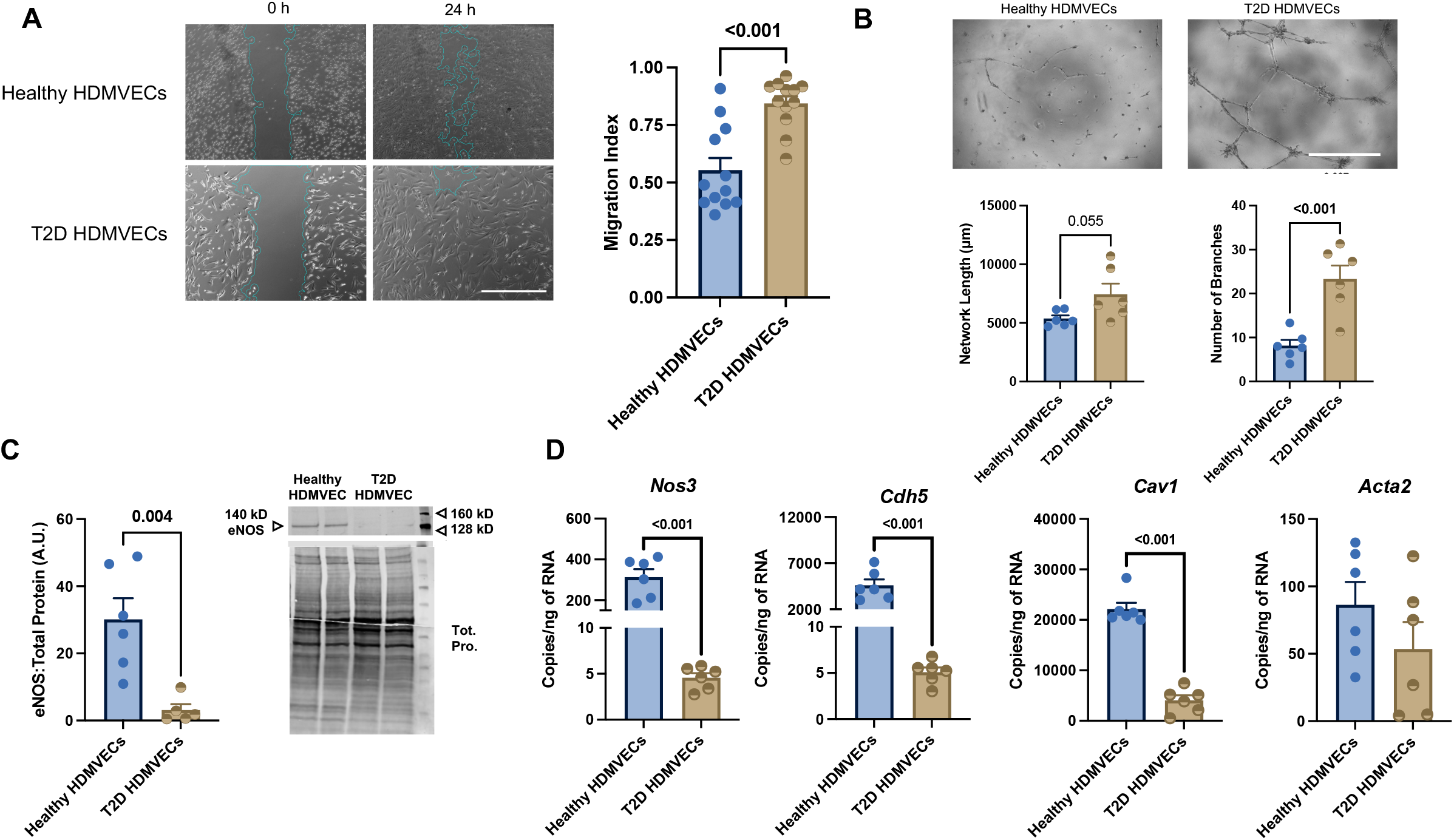
Endothelial cell migration and protein/gene expression in Healthy-HDMVECs and T2D-HDMVECs. **A**) Representative images for the scratch assay in Healthy-HDMVECs (n = 12) and T2D-HDMVECs (n = 12) are shown at baseline (0 hours) and 24 hours after the scratch. Scale bars represent 1000 µm. Image analysis was obtained for all 24 replicates, and the migration index was calculated. **B**) Representative images for the tube formation assay are shown for Healthy-HDMVECs (n = 6) and T2D-HDMVECs (n = 6) 24 hours after re-plating on Matrigel. Scale bars represent 1000 µm. Network length and the number of branches were quantified. **C**) eNOS protein expression in Healthy-HDMVECs (n = 6) and T2D-HDMVECs (n = 5) was detected via Western blotting and normalized to total protein. **D**) Gene expression of Nos3, Cdh5, Cav1 (assessed via Mann-Whitney U test), and Acta2 was measured via ddPCR in Healthy-HDMVECs (n = 6) and T2D-HDMVECs (n = 6). Group differences were determined via independent t-tests, or Mann-Whitney U tests if denoted. Data are expressed as mean ± SEM.

### Effects of Healthy-serum and T2D-serum on Healthy-HDMVECs and T2D-HDMVECs

The migration indices were not statistically significant between conditions (**Fig. 3A**); however, exposure to both Healthy-serum and T2D-serum blunted tube formation in Healthy-HDMVECs compared to the serum-free control condition (**Fig. 3B**). For example, exposure to Healthy-serum and T2D-serum reduced network length by 51% (*p* = 0.082) and 70% (*p* = 0.011), respectively, while the number of branches was reduced by 64% (*p* = 0.117) and 79%, respectively (*p* = 0.038; **Fig. 3B**). The patterns of nitrite production — a biomarker for nitric oxide (NO) secretion — from Healthy-HDMVECs were consistent with the patterns observed for tube formation. For example, exposure to Healthy-serum (−169%, p = 0.052) and T2D-serum (−175%, p = 0.043) blunted nitrite production compared to the serum-free control condition (**Fig. 3C**). These effects on tube formation and nitrite were not observed in T2D-HDMVECs, nor were there differential effects between the exposures to Healthy-serum and T2D-serum (**Fig. 3B** and **Fig. 3C**).

**Fig. 3.**
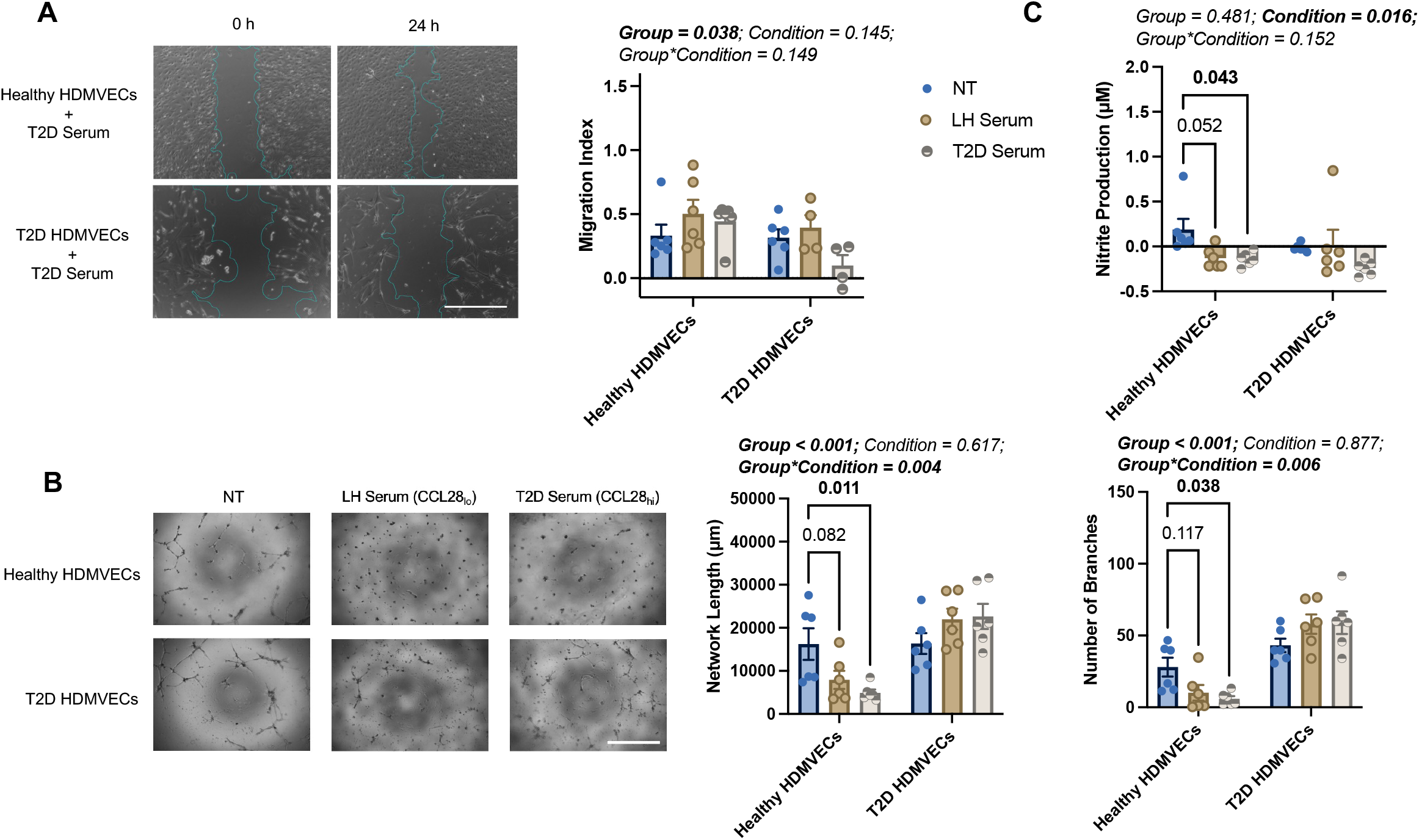
Effects of human serum on cell migration, tube formation, and nitrite production in Healthy-HDMVECs and T2D-HDMVECs. **A**) Representative images for the scratch assay following exposure to serum-free medium (Healthy-HDMVECs = 6; T2D-HDMVECs = 6), Healthy-serum (Healthy-HDMVECs = 6; T2D-HDMVECs = 4), and T2D-serum (Healthy-HDMVECs = 6; T2D-HDMVECs = 4) are shown at baseline (0 hours) and 24 hours after the scratch. Scale bars represent 1000 µm. Image analysis was obtained for 32 of the 36 replicates, and the migration index was calculated. **B**) Representative images for the tube formation assay following exposure to serum-free medium (Healthy-HDMVECs = 6; T2D-HDMVECs = 6), Healthy-serum (Healthy-HDMVECs = 6; T2D-HDMVECs = 6), and T2D-serum (Healthy-HDMVECs = 6; T2D-HDMVECs = 6) are shown, 24 hours after replating on Matrigel. Scale bars represent 1000 µm. Network length and the number of branches were quantified. **C**) Nitrite production during the 24-hour exposures was calculated. Two-way ANOVAs were used to identify significant differences between conditions and treatments. Šídák’s tests were used to assess pairwise multiple comparisons. Data are expressed as mean ± SEM.

Consistent with the decrease in tube formation and nitrite production, the expression of genes involved in the Notch signaling pathway (*Hes1, Hey1, Dll4, Jag1, Notch1, Notch2*, and *Notch3*) and angiogenesis (*Vegfr2*) was markedly lower in Healthy-HDMVECs following exposure to either Healthy-serum or T2D-serum, when compared to serum-free medium (**Fig. 4**). For example, Healthy-serum and T2D-serum treatments reduced *Hes1* by 99% (*p* < 0.001) and 92% (*p* < 0.001), *Dll4* by 84% (*p* < 0.001) and 65% (*p* < 0.001), *Notch1* by 58% (*p* < 0.001) and 64% (*p* < 0.001), *Notch2* by 90% (*p* = 0.003) and 60% (*p* = 0.068), *Notch3* by 80% (*p* < 0.001) and 99% (*p* < 0.001), and *Vegfr2* by 93% (*p* < 0.001) and 77% (*p* < 0.001), respectively. In T2D-HDMVECs, however, the expression of many of these genes (*Hes1*, P=0.061; *Hey1*, P=0.022; *Jag1*, P<0.001; *Notch2*, P=0.002; and *Vegfr2*, P=0.028) was greater following exposure to Healthy-serum, whereas their expression was unaffected by T2D-serum.

**Fig. 4.**
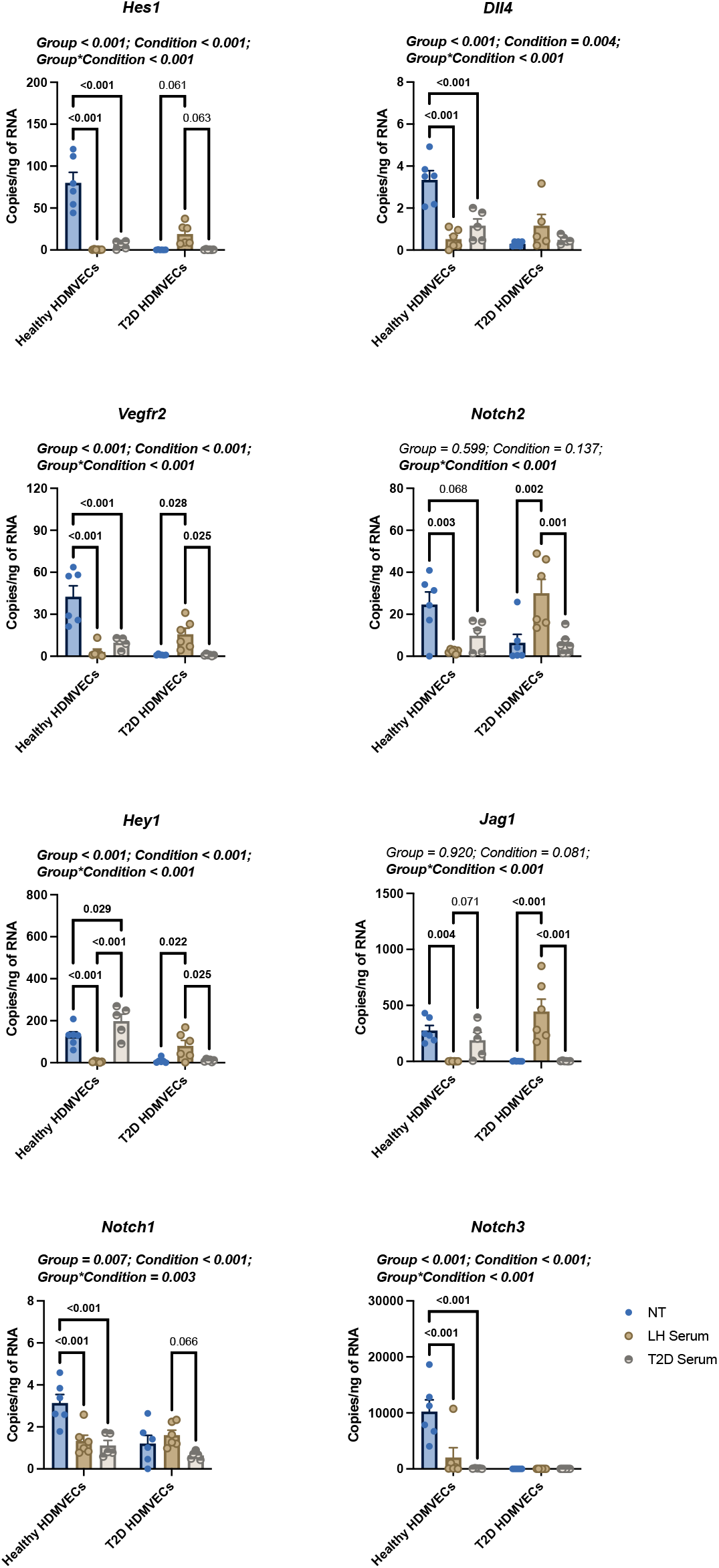
Effects of human serum on gene expression in Healthy-HDMVECs and T2D-HDMVECs. The gene expression of *Hes1, Hey1, Dll4, Jag1, Vegfr2, Notch1, Notch2*, and *Notch3* was measured via ddPCR in cell supernatants following exposure to serum-free medium (Healthy-HDMVECs n = 6; T2D-HDMVECs n = 6), Healthy-serum (Healthy-HDMVECs n = 5; T2D-HDMVECs n = 6), and T2D-serum (Healthy-HDMVECs n = 6; T2D-HDMVECs n = 6). Two-way ANOVAs were used to identify significant differences between conditions and treatments. Šídák’s tests were used to assess pairwise multiple comparisons. Data are expressed as mean ± SEM

## Discussion

We hypothesized that human dermal microvascular endothelial cells (HDMVECs) from individuals with T2D would exhibit impaired endothelial function. Yet, in Experiment 1 (**Fig. 2**), our data paradoxically show that T2D-HDMVECs have enhanced wound-healing capacity (i.e., elevated migration index) and angiogenic potential (i.e., longer and more branched tube networks) compared to healthy controls. Despite these seemingly advantageous traits, T2D-HDMVECs also exhibited elongated and disorganized morphology, nearly undetectable eNOS expression, and markedly reduced expression of endothelial genes (*Cdh5* and *Cav1*), while mesenchymal gene expression (*Acta2*) remained unchanged. Because EndMT is characterized by mesenchymal morphology, increased cell motility, and decreased expression of canonical endothelial genes—which ultimately leads to aberrant endothelial function [15, 16]—these findings imply that T2D-HDMVECs display an EndMT phenotype and engage in pathological angiogenesis when damaged. Such a loss of endothelial identity would be detrimental to wound healing in T2D.

We further hypothesized that serum from individuals with T2D would compromise endothelial function in HDMVECs from healthy donors. Indeed, T2D-serum reduced both tube formation (angiogenesis) and nitrite (NO) production in Healthy-HDMVECs, suggesting that at least part of the endothelial dysfunction in T2D could stem from exposure to circulating factors. However, Healthy-serum produced comparable effects. Interestingly, neither Healthy-nor T2D-serum affected migration, angiogenesis, or nitrite production in T2D-HDMVECs, suggesting that T2D-HDMVECs are unresponsive (or resistant) to these circulating factors. Moreover, the similar effects observed with both Healthy- and T2D-serum on Healthy-HDMVECs indicate that the diabetic state does not substantially alter the circulating factors that regulate endothelial migration or angiogenesis. Alternatively, these data could also indicate that donor serum caused immunogenic responses that were not tested for, which is a limitation to this type of experimental design. Future experiments would be needed to elucidate the causative factors for the observed phenotypes.

We also explored whether Notch-dependent mechanisms might underlie the effects of T2D-serum. Under serum-free conditions, Healthy-HDMVECs showed higher levels of Notch/angiogenic gene expression, whereas adding serum (from either healthy or T2D donors) suppressed these genes in Healthy-HDMVECs. By contrast, T2D-HDMVECs displayed a different pattern: they upregulated several Notch-related genes and Vegfr2 after exposure to Healthy-serum, but not T2D-serum. These nuances emphasize the importance of Notch signaling.

### Baseline differences between Healthy-HDMVECs and T2D-HDMVECs

Although T2D-HDMVECs originate from a diabetic milieu, they paradoxically demonstrated ∼52% higher scratch closure (i.e., migration) and more robust tube formation compared with Healthy-HDMVECs—suggestive of a “hyperangiogenic” phenotype. Under normal conditions, eNOS-derived NO is vital for healthy vascular function (e.g., vasodilation and controlled angiogenesis). Thus, the near-absence of eNOS in T2D-HDMVECs implies reliance on alternative, and likely dysregulated, pro-angiogenic signals. Both *Cdh5* (VE-Cadherin) and *Cav1* (Caveolin-1) are essential for preserving endothelial integrity and function (including eNOS regulation); their downregulation in T2D-HDMVECs underscores a dysfunctional endothelial state prone to forming aberrant or unstable vessels. Meanwhile, the lack of change in *Acta2* indicates that T2D-HDMVECs are not transitioning toward a smooth muscle-like phenotype. Overall, T2D-HDMVECs exhibit paradoxically heightened angiogenic/migratory activity coupled with a dysfunctional eNOS/NO axis — likely reflecting a pathological form of angiogenesis commonly associated with diabetes [8, 9].

### Responses of Healthy-HDMVECs and T2D-HDMVECs to Healthy-Serum vs. T2D-Serum

In Healthy-HDMVECs, exposure to either healthy- or T2D-serum diminished tube formation relative to serum-free conditions. Nitrite (a marker for NO) production also declined, suggesting that serum factors may transiently suppress eNOS expression or Notch signaling in normally quiescent cells. Notably, there were no major differences between Healthy- and T2D-serum in their effects on Healthy-HDMVECs, indicating that both serum types exert a similar inhibitory influence on NO-dependent angiogenic pathways in healthy cells. In contrast, T2D-HDMVECs did not exhibit a substantial decrease in tube formation or nitrite production after exposure to either serum type, suggesting that they are resistant to the inhibitory effects seen in Healthy-HDMVECs. Their low eNOS (*Nos3*) expression may render T2D-HDMVECs less sensitive to external modulators of eNOS, allowing them to remain in a hyperangiogenic state regardless of serum type.

Gene expression analyses provided additional insights. In Healthy-HDMVECs, both healthy- and T2D-serum suppressed Notch pathway genes (*Hes1, Hey1, Dll4, Jag1, Notch1/2/3*) and *Vegfr2*, mirroring the reductions in tube formation and NO production. Conversely, in T2D-HDMVECs, several of these genes (*Jag1, Notch2, Vegfr2*) were upregulated only in response to Healthy-serum, suggesting that T2D-HDMVECs can benefit from restorative factors present in Healthy-serum — factors apparently absent or neutralized in T2D-serum.

This is the first study examining the impact of human serum on primary human dermal endothelial cells. Other T2D cell types show similar “resistance” to stimuli: for example, primary monocytes from individuals with T2D exhibit blunted TNF-α secretion and reduced CD11b and TLR4 expression in response to LPS, compared to monocytes from healthy donors [37]. Such parallels are relevant because monocytes/macrophages and endothelial cells share a common hemangioblast lineage [38], highlighting the intertwined nature of the hematopoietic and vascular systems. Much like insulin resistance — a hallmark of T2D — the vasculature and immune cells also appear resistant to certain circulating cues. The specific identity and relevance of these factors warrant further investigation.

### Possible implications

T2D-HDMVECs present a dysfunctional, hyperangiogenic phenotype: they migrate and form tubes more readily but have markedly reduced eNOS expression, suggesting a shift from protective NO-dependent mechanisms toward potentially pathological remodeling. Healthy-HDMVECs, meanwhile, exhibit a more regulated angiogenic capacity and a clear responsiveness to serum factors, which transiently suppress eNOS/NO and Notch signaling. Notably, both healthy- and T2D-serum had similar dampening effects on healthy cells; however, T2D-serum failed to upregulate certain *Notch* and *Vegfr2* genes in T2D-HDMVECs, whereas healthy serum did. This discrepancy indicates that T2D-serum may lack (or contain antagonists of) the beneficial signals that partly restore normal gene expression in T2D cells.

The synchronous decrease (in healthy cells) or partial upregulation (in T2D cells exposed to healthy serum) in Notch targets (*Hes1, Hey1, Dll4, Jag1, Notch1/2/3*) and *Vegfr2* strongly supports a pivotal role for the Notch pathway in orchestrating angiogenesis and NO biology in these cells. Overall, T2D endothelial cells appear “reprogrammed”, displaying a heightened baseline migratory/angiogenic drive, decreased reliance on NO, and reduced responsiveness to serum-derived inhibitory signals. Healthy endothelial cells, on the other hand, depend on eNOS/NO and Notch for controlled angiogenesis and are more susceptible to serum-induced suppression of these pathways. Notably, T2D endothelial cells and T2D serum both contribute to an abnormal gene expression profile, but T2D cells can partially regain normal Notch/angiogenesis signaling when exposed to healthy serum. This underscores the interplay between the cells’ intrinsic state and the extrinsic milieu (serum), indicating that impaired wound healing in T2D arises from a convergence of “intrinsically altered” endothelial function and suboptimal circulating factors.

### Future research

Notch signaling — typically via Notch 1/4 — is known to regulate wound healing [21, 22] and studies show that high glucose can activate Notch1 [39]. In rodent models of diabetes, Notch activation impairs wound healing, while chemical and genetic inhibition of Notch signaling improves it [39]. Our data extend these observations into clinical models, reinforcing Notch inhibition as a potential therapeutic strategy for accelerated healing of diabetic foot ulcers [22].

Our prior work established that CCL28/CCR10 signaling modulates eNOS activity [18, 19] and is required for wound repair in a T2D mouse model [19, 20]. Given the marked downregulation of eNOS in T2D-HDMVECs noted here, disrupting CCL28/CCR10/eNOS interactions may be a viable therapeutic strategy for non-healing wounds, including diabetic foot ulcers. There is a high clinical need for novel therapeutic targets, especially because the last approved drug for wound healing — becaplermin — occurred in 1997.

This study also suggests that T2D serum harbors (or lacks) factors with a profound influence on endothelial processes essential for wound repair (migration, angiogenesis, and NO signaling). Identifying these “bioactive” components—whether proteins, lipids, nucleic acids, or other molecules— is a critical next step [40]. Such knowledge could guide the development of targeted therapies, including mimetics or inhibitors of these factors, to reestablish normal endothelial function in T2D. Therapeutic approaches might involve supplementing beneficial factors absent from T2D serum or blocking pathological signals that drive dysregulated angiogenesis and inadequate wound healing.

Because T2D wound healing is influenced by various systemic factors (e.g., inflammation, metabolic dysfunction) [2, 3, 41], understanding inter-individual variations in these circulating components could enable more personalized treatments. Ultimately, pinpointing the nature and function of these molecules will be essential for designing improved therapies for individuals with type 2 diabetes.

### Limitations

First, we selected the 24-hour exposure period based on unpublished pilot data; without a complete time course, it remains possible that T2D-HDMVECs might respond differently over longer exposures or at varying serum concentrations. Second, although primary human endothelial cells provide better clinical translatability than rodent-derived cell lines, the varied genetic and environmental factors that contribute to T2D may limit the generalizability of our findings. Third, recent single-cell studies have identified endothelial subtypes expressing signatures for phagocytosis/scavenging, antigen presentation, and immune recruitment [42]. These cells can express receptors for pathogen-associated molecular patterns and major histocompatibility complex class II (MHC-II), meaning that exposure to foreign elements in serum could trigger immune-related pathways and confound our interpretation of serum-induced effects. Further, our *in vitro* model using endothelial monolayers does not perfectly recapitulate *in vivo* vasculature. Finally, the healthy individuals were not age, sex, or BMI matched between groups; however, we contended that any deviation from the healthy state is indicative of T2D pathophysiology. Future work employing 3D cultures or co-cultures (with fibroblasts, immune cells, etc.) would yield a more physiologically relevant model of the wound microenvironment. Despite these limitations, using clinically relevant primary cell lines produced novel findings that advance understanding of wound healing processes.

### Conclusion

Impaired wound healing in T2D arises from a convergence of intrinsic endothelial reprogramming and dysregulated extrinsic cues. Although T2D-HDMVECs displayed unexpectedly hyperangiogenic features, they remained largely unresponsive to putative serum-derived signals, highlighting disrupted eNOS and Notch signaling. By contrast, healthy endothelial cells showed robust responses to both healthy and T2D serum, underscoring the importance of circulating factors for regulating angiogenesis. These observations suggest that targeting the intrinsic cellular pathways (e.g., Cav1, Notch-1, eNOS) and the extrinsic humoral environment (e.g., supplementation or blockade of specific circulating factors such as CCL28) could improve wound healing in individuals with T2D. Given that diabetic foot ulcers pose a major clinical challenge, and because the current standard of care is often applied heterogeneously and may be inadequate [2], our results provide a foundation for future strategies that optimize patient care.

## Acknowledgements

We acknowledge Michelle Pieper for her technical expertise in figure editing.

## Author contributions (CRediT)

Conceptualization: JS, SG, RM, AL, JH

Data curation: JS, CM, PV, JK, AA

Formal analysis: JS, JH

Funding acquisition: JH, CH, MM, BS

Investigation: JS, CM, PV, JK, AA

Methodology: JS, CM, PV, JK, AA, SG, RM, AL, JH

Project administration: JS, AL, JH

Resources: AL, JH

Software: n/a

Supervision: AL, JH

Validation: SG, RM, TS, AL, JH

Visualization: JS, SG, RM, TS, AL, JH,

Writing – original draft: all authors

Writing – review and editing: all authors

## Data availability statement

Data is available from the Corresponding Author.

